# AAV-mediated genome editing is influenced by the formation of R-loops

**DOI:** 10.1101/2024.05.07.592855

**Authors:** Francesco Puzzo, Magdalena P. Crossley, Aranyak Goswami, Feijie Zhang, Katja Pekrun, Jada L. Garzon, Karlene A. Cimprich, Mark A. Kay

**Author notes:** These authors contributed equally to this work (Francesco Puzzo, Magdalena P. Crossley).

## Abstract

Recombinant adeno-associated viral vectors (rAAV) hold an intrinsic ability to stimulate homologous recombination (AAV-HR) and are the most used in clinical settings for *in vivo* gene therapy. However, rAAVs also integrate throughout the genome. Here, we describe DNA-RNA immunoprecipitation sequencing (DRIP-seq) in murine HEPA1-6 hepatoma cells and whole murine liver to establish the similarities and differences in genomic R-loop formation in a transformed cell line and intact tissue. We show enhanced AAV-HR in mice upon genetic and pharmacological upregulation of R-loops. Selecting the highly expressed *Albumin* gene as a model locus for genome editing in both *in vitro* and *in vivo* experiments showed that the R-loop prone, 3’ end of *Albumin* was efficiently edited by AAV-HR, whereas the upstream R-loop-deficient region did not result in detectable vector integration. In addition, we found a positive correlation between previously reported off-target rAAV integration sites and R-loop enriched genomic regions. Thus, we conclude that high levels of R-loops, present in highly transcribed genes, promote rAAV vector genome integration. These findings may shed light on potential mechanisms for improving the safety and efficacy of genome editing by modulating R-loops and may enhance our ability to predict regions most susceptible to off-target insertional mutagenesis by rAAV vectors.

## Introduction

Gene therapy holds great promise for treating genetic diseases. Due to their safety and efficacy, recombinant adeno-associated viruses (rAAV, here after just AAV) are the vectors of choice for in vivo gene transfer in clinical settings^1,2^ and have been recently approved for treatment of hemophilia A and B, Leber’s congenital amaurosis, spinal muscular atrophy, and Duchenne muscle dystrophy (https://www.fda.gov/vaccines-blood-biologics/cellular-gene-therapy-products/approved-cellular-and-gene-therapy-products).

The AAV genome forms extrachromosomal (episomal) concatemers in the cell nucleus upon viral transduction^3^. Therefore, treatment of pediatric subjects would be hampered due to cell division during organ development resulting in loss of the episomal vector^4^. To overcome this issue, we have devised a nuclease-free genome editing tool (AAV-HR) which targets the 3′ end of *Albumin (Alb)* using homology arms surrounding a self-cleaving peptide (P2A) and a promoterless gene of interest^5^. This technology takes advantage of 1) homologous recombination occurring in the vector-transduced cells to permanently integrate the AAV cassette into the genome without the use of nucleases, which could induce harmful off-target cleavage and DNA double strand breaks^6^, and 2) high expression and secretion of albumin, which in turn leads to high cellular expression and secretion of the gene of interest upon AAV-HR transduction.

AAVs have been reported to naturally integrate at very low rates into the host genome upon cell transduction^7,8^. Recombinant AAVs contain inverted terminal repeats (ITR), the only viral sequence present in the AAV cassette, which possess recombinogenic properties that may drive viral integration^9,10^.

Interestingly, transcriptionally active genomic regions have been identified as one of the main hotspots for AAV-HR^11^ and vector integration^12,13^ yet transcription alone is not the only parameter dictating these events^11^. Moreover, inhibition of non-homologous end-joining (NHEJ) DNA repair complexes has been shown to enhance AAV-HR^14^, while knocking down proteins involved in homologous recombination (HR) repair resulted in the repression of AAV-HR^15^. Nevertheless, the mechanisms underlying AAV-HR and AAV integration events in the genome still remain elusive.

We recently identified the Fanconi anemia complementation group M (FANCM) as one factor that modulates AAV-HR in mammalian cells. Specifically, loss of FANCM increased the efficiency of AAV-HR in human cells^16^. FANCM is also known to regulate the resolution of RNA-DNA hybrid-containing structures called R-loops^17^. R-loops form on the genome during transcription when the newly synthesized RNA molecule anneals to its template DNA and in turn displaces the other DNA strand. R-loops have multiple physiological roles in cells, promoting efficient transcription initiation and termination^18^. Conversely, R-loops can also have potentially destabilizing effects on the underlying DNA molecule. R-loops have been linked to increased genome instability and are associated with the phenomena of transcription-associated recombination (TAR) and transcription-associated mutagenesis (TAM), whereby highly transcribed genes exhibit increased genomic instability^19^. R-loops frequently form at highly expressed genes in mammalian cells^20^ and can be a source of DNA damage - for example by exacerbating transcription-replication conflicts^21^, or due to the processing of DNA repair nucleases which can act on R-loops^22^. RNA-DNA hybrids have also been implicated in directing various aspects of DNA double-strand break repair pathways^23,24^. For example, within transcriptionally active regions, transcription stimulates homologous recombination in an R-loop-dependent manner in cells^25^. However, links between R-loop formation and AAV-associated recombination, and how this might influence the outcomes of gene therapy, have not hitherto been reported. Here, we use genetic and pharmacological approaches to modulate R-loops and show that R-loop levels influence AAV-HR and AAV integration. We mapped R-loops genome-wide in cells and in mouse tissue, revealing how high levels of R-loops at the 3’ end of a highly expressed gene, such as *Alb*, potentially promote AAV-HR. Considering these findings, we anticipate that R-loop formation is an important co-transcriptional process that modulates AAV vector integration and could have implications for gene-based therapeutics.

## Results

### HEPA 1-6 murine cells display higher levels of genomic R-loops compared to mouse liver tissue

We previously demonstrated successful nuclease-free genome editing by liver-mediated AAV gene transfer and developed AAV-HR^5^. To probe how R-loop formation may affect AAV-HR and associated integration events, we mapped R-loops genome-wide using DNA-RNA immunoprecipitation with sequencing (DRIP-seq) in whole mouse liver and, for comparison, in liver-derived murine hepatoma cells, HEPA 1-6 (Fig. 1a). We optimized the DRIP procedure for snap-frozen mouse tissues, homogenizing mouse liver prior to extracting intact genomic DNA for immunoprecipitation (Fig.1b and Materials and Methods). Notably, we validated this approach by DRIP-qPCR analysis of snap-frozen mouse brain tissue, providing proof-of-principle that our approach expands the utility of DRIP-seq to primary tissues that can be frozen (Extended Data Fig. 1a).

**Figure 1:**
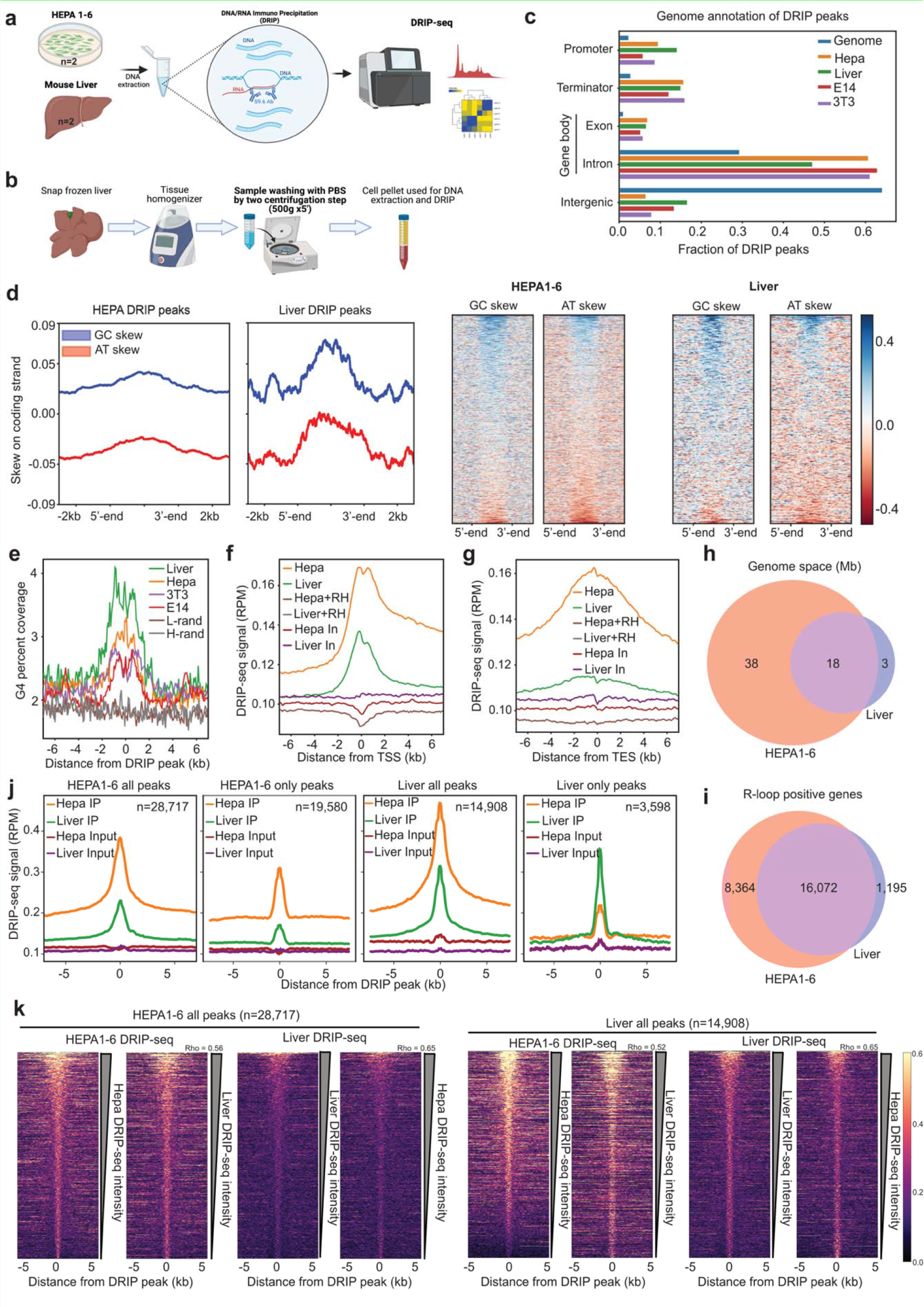
DRIP-seq analysis. **a.** Schematic of the DRIP-seq workflow for HEPA 1-6 and murine liver. Two biological replicates for each sample were sequenced. **b.** Schematic of whole tissue preparation for DRIP- seq. **c.** Fractions of DRIP peaks overlapping murine genomic features from HEPA1-6 (Hepa), liver, E14 embryonic stem cells (E14) and 3T3 fibroblast cells (3T3). **d.** Left, aggregate plots of mean GC (blue) and AT (red) skew across the coding strand of DRIP peaks, including 2 kb flanking the 5’- and 3’-ends. Bands represent 95% CI of mean skew signal. Right, heatmaps showing skew across all individual peaks plotted on the left, ranked by magnitude of skew. Scale bar shows skew values. **e.** Aggregate plots around the center of DRIP peaks showing the G- quadruplex motif density. **f.** Aggregate plot of DRIP-seq signal around the TSS of mouse genes, showing signal from IP, inputs (In) and RNaseH-treated (+RH) samples. **g.** As for f, but around the TES of mouse genes. **h.** Venn diagram of genome areas (in megabases; Mb) occupied by peaks called from DRIP IP vs input in HEPA1-6 and liver tissue. **i.** Venn diagram of R-loop positive genes in HEPA1-6 and liver tissue. **j)** Aggregate plots showing DRIP-seq signal around the center of different sets of DRIP peaks from HEPA1-6 and liver. IP and input samples are shown. **h.** Heatmaps showing the DRIP-seq signal around the center of peaks called in HEPA1-6 (left) or liver (right). Heatmaps are ranked by DRIP-seq signal strength within 3kb of the peak center. Correlation coefficients (Spearman’s Rho) are shown.

We first compared our DRIP-seq data with those previously obtained using other murine cell lines^26^. HEPA 1-6 and liver R-loop forming sites were similarly found mostly in actively transcribed regions such as exons, introns, gene promoters and terminators, whereas they were underrepresented in intergenic regions^27^ (Fig. 1c). HEPA1-6 and liver R-loop forming regions clearly exhibited a pattern of GC-skew (asymmetry in G content between strands) over AT-skew (asymmetry in A content between strands) (Fig. 1d), consistent with DRIP peaks in other mouse cell lines (Extended Data Fig. 1B)^26,28^. We also observed similar nucleotide content across all murine DRIP peaks (Extended Data Fig. 1c, d). Since regions of high G-skew are associated with secondary structure formation such as G-quadruplexes^26^, we examined the propensity of secondary structure formation across DRIP peaks in mouse cells, using non-B form DNA secondary structure predictions^29^. We found strong enrichment for G4-forming motifs (Fig. 1e), but not other non-B DNA secondary structures, at mouse DRIP peaks, and particularly within peaks from HEPA1-6 and liver (Extended Data Fig. 1e). This may reflect a higher proportion of peaks in HEPA1-6 and liver forming in gene promoters (Fig. 1c), where G-quadruplexes are enriched^30^.

Subsequently, we compared RNA-DNA hybrid formation genome-wide between HEPA1-6 and liver tissue. We observed robust enrichment of DRIP-seq signal above input at well characterized R-forming genes, including *Apoe* and *Actb* (Extended Data Fig. 2a), and because these signals were RNaseH-sensitive, it indicated that the DRIP-seq captured bona-fide RNA-DNA hybrids. We next asked how the DRIP-seq hybrid signals correlated with known features of the genome. Across promoters, DRIP-seq signal peaked sharply within 1 kb of the transcription start site (TSS) and extended downstream of the TSS into the gene body (Fig. 1f), consistent with previous reports of R-loops forming robustly across the first exon^31^. At the end of genes, we observed a broader DRIP-seq profile spanning 4 kb each side of the transcription end site (TES) (Fig. 1g), consistent with DRIP-seq capturing R-loops that form at gene terminators upon RNAPII pausing as well as those generated by transcriptional read through beyond the TES. At both TSS and TES regions, the DRIP-seq signals from HEPA1-6 were higher than liver, and in all cases were verified by their RNaseH-sensitivity.

We then performed peak calling against matched input samples. We identified more R-loop-containing sites in HEPA1-6 compared to liver (28,717 peaks in HEPA1-6 versus 14,908 in liver) and these occupied a greater area of the genome (56.4 Mb in HEPA1-6 versus 21.6 Mb in liver) (Fig. 1h). DRIP peaks overlapped with 24,436 genes in HEPA1-6 and 17,267 genes in liver, with most R-loop-positive genes (16,072) common to both cell types (Fig. 1i). HEPA1-6 DRIP-seq signal was higher than liver in common R-loop sites (Fig. 1j) and the DRIP-seq signal strength from both HEPA1-6 and liver correlated positively with each other (Fig. 1k, Extended Data Fig. 2b). These data suggest that DRIP-seq signal was highly congruent in both HEPA1-6 and liver, but that the overall liver DRIP-seq signal was lower at most R-loop forming sites than HEPA1-6. Notably, DRIP-seq signal from the liver was still detectable above input within HEPA1-6 unique peaks (Fig. 1k), but below the threshold of peak calling. Harsher conditions of DNA extraction required for liver tissue may have reduced the overall DRIP-seq enrichment compared to HEPA1-6.

While the peaks that were called uniquely in HEPA1-6 were mostly genic, those called uniquely in liver were mostly intergenic (Extended Data Fig. 2c) and the genic peaks were enriched within long genes (Extended Data Fig. 2d), reflecting tissue-specific R-loops forming in some genes and regulatory regions. In agreement with this, the degree of DRIP-seq signal from HEPA1-6 and liver did not correlate well within these liver-specific sites (Extended Data Fig. 2e), potentially indicating that a subset of R-loops form in a tissue-specific manner but are absent or altered in transformed cells (HEPA1-6).

The nucleotide content of HEPA1-6 and liver DRIP peaks were similar overall, with those peaks identified only in the liver exhibiting a lower GC content than the other peaks (Extended Data Fig. 2f). We did not observe large differences in peak sizes - both HEPA1-6 and liver had a broad distribution with a median length of 1.1 kb (HEPA1-6) and 0.9 kb (liver) and interquartile ranges of 0.6-2.3 kb (HEPA1-6) and 0.4-1.7 kb (liver) (Extended Data Fig. 2g, h). Therefore, the distribution, relative abundance and nucleotide properties of R-loops are largely conserved between transformed HEPA1-6 cells and primary liver tissue. We identified several R-loop sites with unique properties in the liver, which may reflect biological differences between R-loop formation between transformed cell lines and primary tissue.

### DRIP-seq and RNA-seq data correlation studies in liver and HEPA 1-6

To determine how the genomic R-loop landscape is modulated by gene expression, we analyzed RNA-seq datasets from HEPA 1-6 and murine liver. We found 7,456 genes upregulated and 8,086 downregulated in liver compared to HEPA1-6 (Fig. 2a), with HEPA1-6 exhibiting a higher transcriptional activity (Fig. 2b). Overall, we found no correlation between HEPA1-6 and liver RNA-seq (Fig. 2c), emphasizing the profoundly different transcriptome between transformed cells and primary tissue. Consistent with higher cell proliferation levels, we found upregulation of genes involved in DNA replication, splicing and ribosome function in HEPA1-6, and down regulation of quiescence genes (Fig. 2d and Supplementary Table 1).

**Figure 2:**
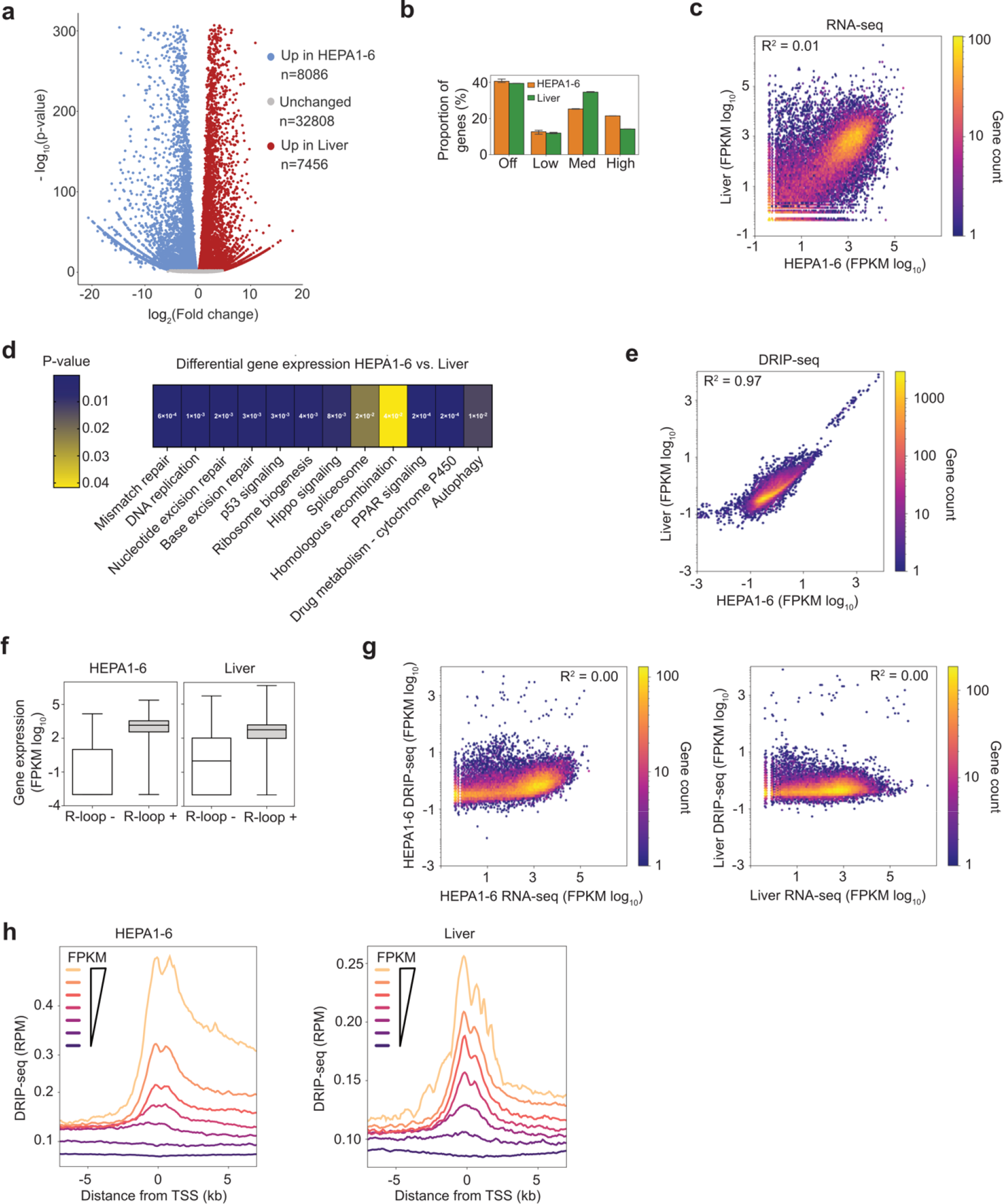
RNA-seq analysis and comparison with DRIP-seq in HEPA1-6 and liver. a. Volcano plot showing downregulated (blue) and upregulated (red) genes in HEPA1-6 vs. Liver, gene numbers a shown. **b.** Gene expression levels in HEPA1-6 and liver. Expression level cutoffs are from EMBL-EBI Expression Atlas (https://www.ebi.ac.uk/gxa/): off (<0.5 FPKM), low (between 0.5 to 10 FPKM), medium (between 11 to 1000 FPKM), high (above 1000 FPKM). **c**. Scatter plot showing RNA-seq FPKM values for 65,440 mouse genes, Pearson’s correlation coefficient is shown. **d**. Heatmap graph showing differential expression analysis between HEPA1-6 and murine liver RNA-seq data (HEPA1-6 vs. Liver; n=3). **e.** Scatter plot showing DRIP-seq signal at 65,440 mouse genes, Pearson’s correlation coefficient is shown. **f.** Gene expression level (FPKM) of genes overlapping (R-loop positive) or not (R-loop negative) with DRIP-seq peaks in HEPA1-6 and liver. Box: 25th and 75th percentiles; central line: median. **g.** Scatter plot showing DRIP-seq signal vs RNA-seq signal at 65,440 mouse genes for HEPA1-6 (left) and liver (right), Pearson’s correlation coefficients are shown. **h.** Aggregate plots of DRIP- seq signal around the TSS of mouse genes for HEPA1-6 (left) and liver (right). Mouse genes were sorted by gene expression levels and distributed in octiles of increasing FPKM.

However, there was concordance between HEPA1-6 and liver DRIP-seq signals (Fig. 2e). Since R-loops form co-transcriptionally and are known to be enriched in active genes^32^, we compared RNA-seq with DRIP-seq data in HEPA1-6 and liver datasets. For both cell types, expression levels in genes that contained an R-loop were higher than those without (Fig. 2f). While there was not a strong correlation between RNA-seq and DRIP-seq averaged across whole gene sequences (Fig. 2g), the DRIP-seq signal at the transcription start sites of genes increased with gene expression levels (Fig. 2h), indicating that promoter-associated R-loops were highly dependent on transcriptional output in both cell types.

### In vitro AAV-HR is positively influenced by R-loops at the albumin locus

To evaluate how RNA-DNA hybrids may influence genome editing by liver AAV-mediated gene transfer, we assessed RNA-DNA hybrid formation across the *Alb* locus. Our DRIP-seq data showed that RNA-DNA hybrids were strongly enriched at the 3’ end, which was where AAV-mediated gene editing (AAV-HR) technology was reported to be relatively efficient at integration (Fig. 3a). To validate the DRIP-seq results, we designed three sets of primers targeting distinct regions within the *Alb* gene, including the 3’ end (Fig.3b). Primers for *Actb* were used as a positive control for DRIP (^26^, and Extended Data Fig. 2a). DRIP-qPCR confirmed that the *Alb* locus in both HEPA1-6 and liver harbors differential levels of RNA-DNA hybrids, but both were highest at the 3’ end of the gene (Fig. 3 c, d).

**Figure 3:**
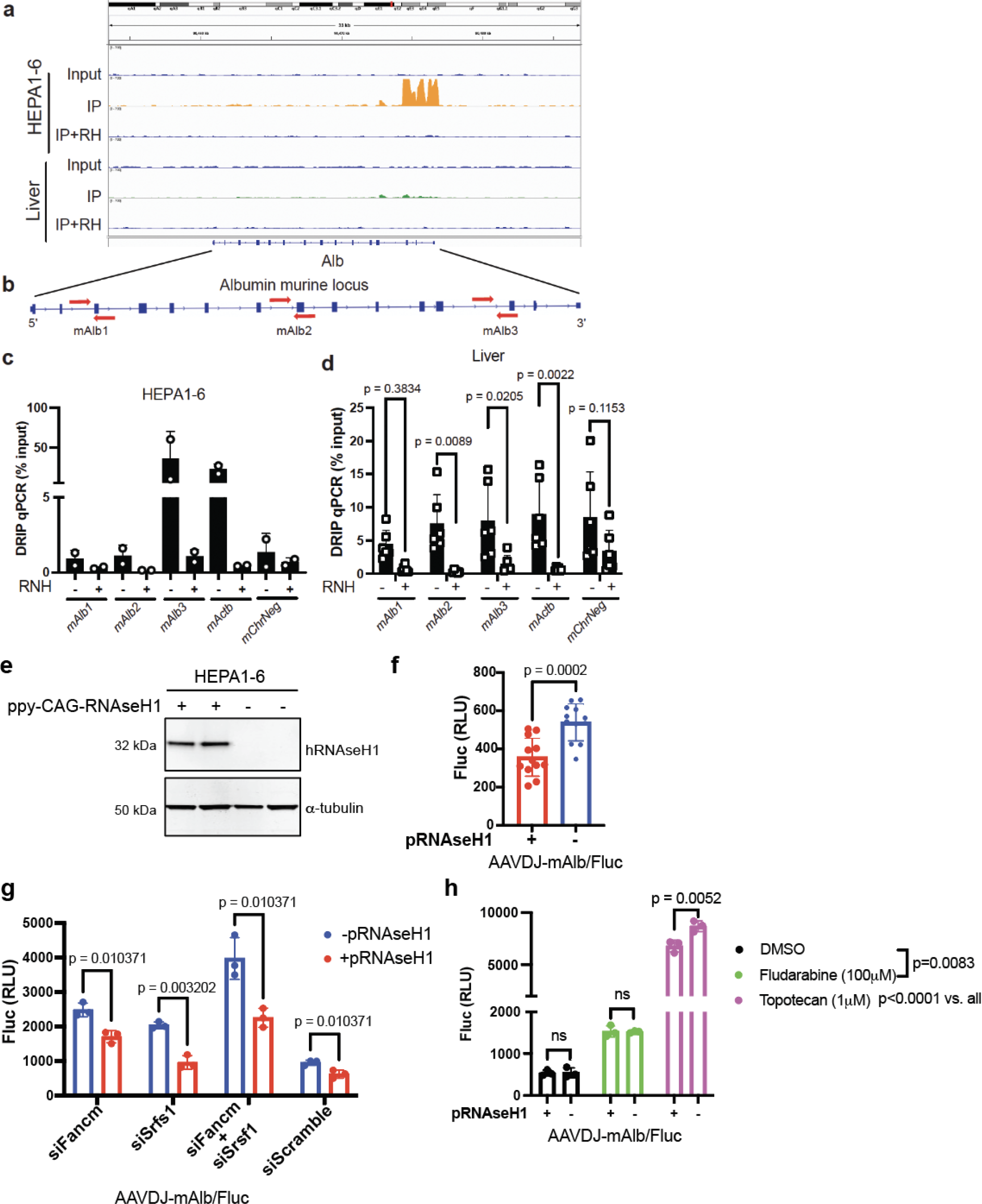
**DRIP-qPCR in liver and HEPA1-6 correlates with DRIP-peaks. a**. Genome browser tracks showing HEPA1-6 and liver DRIP-seq signal at the albumin gen (*Alb)*. Tracks from S9.6 immunoprecipitation (IP), Input and IP with RNaseH treatment (IP+RH) are shown. **b.** Schematic of *Alb* locus and design of primers used for DRIP-qPCR. **c-d.** DRIP- qPCR data in HEPA-1-6 (n=2) (**c**) and liver (n=6) (**d**). RNH is in vitro RNaseH treatment prior to immunoprecipitation. **e.** Western blot showing expression of human RNaseH1 in HEPA1-6 upon transient transfection of the ppy-CAG-RNaseH1 (pRNAseH1) plasmid. **f.** Luciferase activity in HEPA1-6 upon transient transfection of pRNAseH1 plasmid and AAVDJ-mAlb/Fluc transduction (n=12). **g.** Luciferase activity in HEPA1-6 upon transfection of siRNAs against R- loop targeting proteins (siFancm, siSrsf1, or siScramble) and AAVDJ-mAlb/Fluc transduction. Cells were transiently transfected with or without pRNAseH1 plasmid (n=3). **h.** Luciferase activity in HEPA1-6 following treatment with fludarabine, totopotecan or DMSO (negative control) and transduced with AAVDJ-mAlb/Fluc. Cells were transiently transfected with or without pRNAseH1 plasmid (n=3). Statistical analysis: **f.** Unpaired t-test. **d, g, h.** Two-way ANOVA with Sidak’s post hoc analysis. Error bars represent standard deviation of the mean.

To test the hypothesis that R-loops at the 3’ end of the *Alb* gene can positively influence AAV vector genomic integration, we developed a system to easily assess AAV-HR in cells (hereafter referred to as *in vitro*). Based on the previously described AAV-HR vector^5^, we replaced the coding sequence of the human factor IX (hFIX) with the firefly luciferase (Fluc) cDNA surrounded by homology arms targeting the 3’ locus of murine *Alb* (AAV-HR- mAlb/Fluc). We then transduced HEPA 1-6 cells with different multiplicity of infections (MOI) with the AAV-HR-mAlb/Fluc vector and established a vector MOI of 50,000 to use in the following studies (Extended Data Fig. 3a).

Next, to investigate the relationship between R-loop formation and AAV-HR in vitro, we sought to alter R-loop levels by modulating the expression of enzymes involved in R-loop metabolism. To this end, we transiently overexpressed RNaseH1^33^, which degrades the RNA moiety of RNA-DNA hybrids (Fig. 3e). Overexpression of RNAseH1 significantly reduced AAV-HR-mAlb/Fluc mediated integration (Fig. 3f). Next, we used RNA silencing (siRNAs) to knock down other factors implicated in suppressing R-loops (Extended Data Fig. 3b). Knocking down the R-loop resolution factor *Fancm*^16^ resulted in a significant enhancement of AAV-HR- mAlb/Fluc editing in HEPA 1-6 cells, as did knockdown of the splicing factor *Srsf1,* which suppresses R-loop levels^34^ (Extended Data Fig.3c). A slight, but still significant, enhancement was observed when we knockdown *Rnaseh1*^35^. Conversely, depletion of chromatin binding proteins such Sin3a^36^, and two topoisomerases, Top1^34^, and Top3b^37^, had no significant effect on AAV-HR.

RNaseH1 overexpression also significantly reduced AAV-HR-mAlb/Fluc integration upon the silencing of *Fancm* and *Srfs1* gene expression (Fig. 3g), further demonstrating that the removal of R-loops disrupts AAV-HR *in vitro*.

To further confirm these data, we sought to pharmacologically modulate R-loops using topotecan, a topoisomerase1 inhibitor reported to increase R-loop levels^38^. Dose escalation experiments in HEPA 1-6 cells, demonstrated that 1 μM of topotecan significantly increased AAV-HR-mAlb/Fluc integration (Extended Data Fig. 3d), whereas HEPA 1-6 treated with topotecan and in combination with RNaseH1 showed a significant reduction of AAV-HR- mAlb/Fluc expression compared to cells treated with only topotecan (Fig.3h). Conversely, enhanced AAV-HR-mAlb/Fluc expression upon treatment with fludarabine, a small molecule known to enhance AAV-HR^39^, was not reversed by RNaseH1 overexpression, (Fig. 3h) likely because fludarabine does not alter R-loop levels.

Taken together, these results demonstrate that in cells, modulating R-loop formation affects AAV-HR and robust R-loop formation that naturally occurs at the 3’ end of the *Alb* locus promotes AAV-HR.

### AAV-mediated genome integration in vivo is correlated with R-loops levels at the albumin locus

We next investigated whether R-loops could influence AAV-HR *in vivo*. To this end, we treated mice with topotecan, using a dose far below the maximum tolerated^40^ in order to avoid drug-related toxicity (Fig.4a). DRIP-qPCR analysis showed that a systemic dose of 5mg/kg topotecan in vivo increased R-loop levels at the *Alb* locus (Fig.4b).

To assess whether topotecan treatment could enhance AAV-HR *in vivo*, we treated mice with two different doses (5 mg/kg and 10 mg/kg) of topotecan and injected the animals with the previously reported AAV-HR-mAlb/hFIX vector^5^, targeting the 3’ end of *Alb*, which allowed us to test the efficiency of AAV-mediated genome editing by measuring the circulating hFIX in plasma (Fig. 4c). Neither dose of topotecan showed any signs of toxicity in the animals (Extended Data Fig. 4a). Strikingly, we observed a dose-response effect on AAV-HR where both animal cohorts treated with topotecan had significantly higher circulating hFIX compared to mice treated with saline (Fig. 4d). Furthermore, AAV genome vector copy number was higher in the liver of animals treated with the highest dose of topotecan, most likely reflecting elevated AAV integration into the genome (Extended Data Fig. 4b).

**Figure 4:**
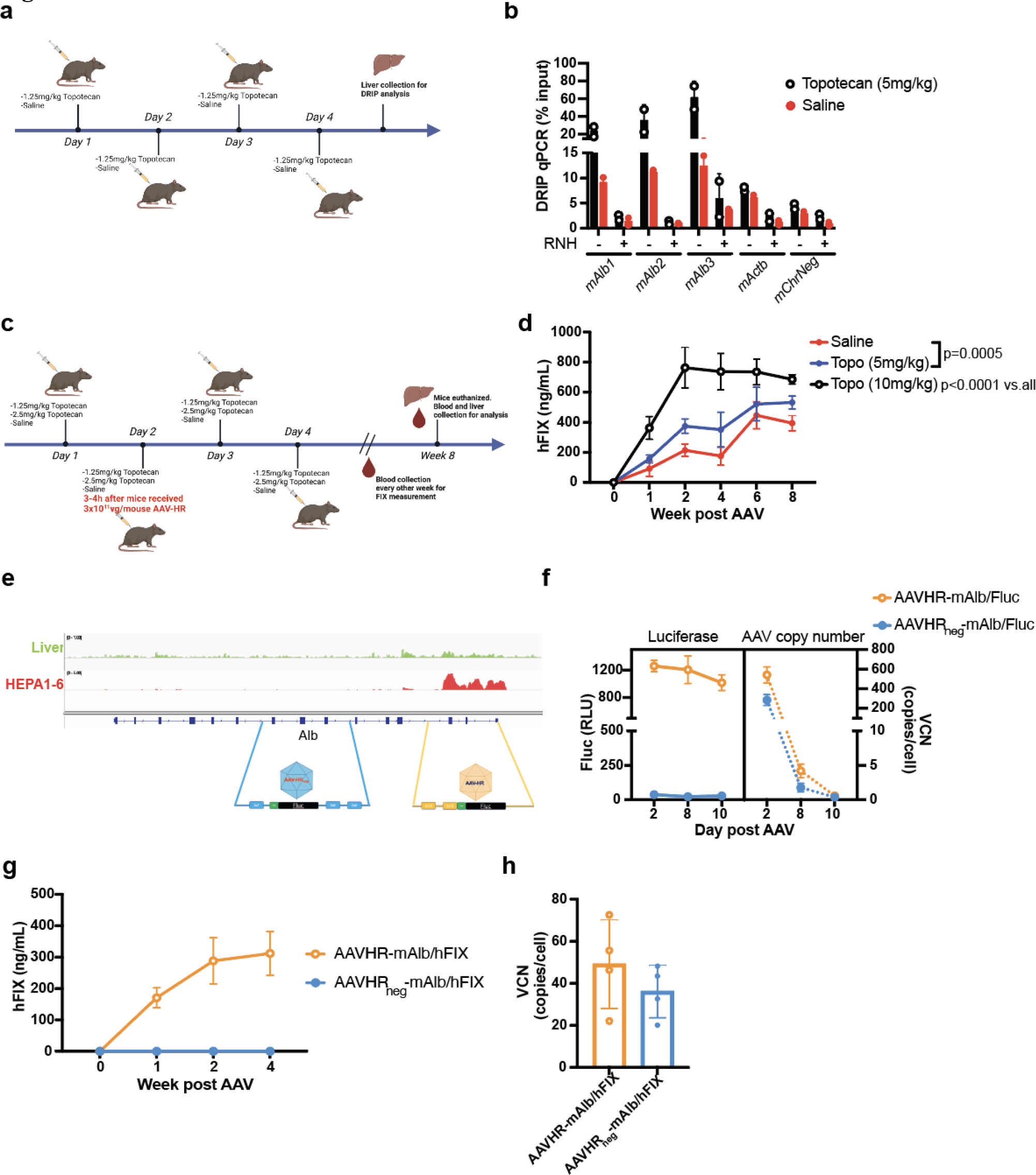
R-loops determine in vitro and in vivo AAV-HR at *Alb* locus. **a.** Schematic representation of topotecan treatment in vivo. **b.** DRIP-qPCR analysis of murine livers upon treatment with 5mg/kg of topotecan (n=2). **c.** Schematic representation of the combination of topotecan treatment and AAV-HR in vivo. **d.** Time course of hFIX circulating levels in mice treated with the combination of AAV-HR-mAlb/hFIX and topotecan (n=3). **e.** Schematic representation of AAV-HR and AAV-HR_neg_ design for genome targeting at the *Alb* locus. **f.** Time course of luciferase activity and AAV vector genome copy number in HEPA1-6 transduced with AAV-HR-mAlb/Fluc and AAV-HR_neg_-mAlb/Fluc (n=3). **g.** Time course of hFIX circulating levels in mice treated with either AAV-HR-mAlb/hFIX or AAV-HR_neg_- mAlb/hFIX (n=4). **h.** AAV vector genome copy number in liver of mice treated with either AAV-HR-mAlb/hFIX or AAV-HR_neg_-mAlb/hFIX (n=4). Statistical analysis: **d.** Two-way ANOVA with Sidak’s post hoc analysis. Error bars represent standard deviation of the mean.

To further investigate the role of R-loops in AAV-mediated genome integration, we made a new AAV-HR vector (AAV-HR_neg_-mAlb/Fluc), targeting a region in the gene body of *Alb* with negligible R-loop formation, hence the nomenclature AAV-HR_neg_ (region negative in R-loops) (Fig. 4e). We designed a series of experiments in which we used two vectors (AAV-HR_neg_ and AAV-HR) targeting the *Alb* locus where the integration sites were within 4 kb of each other in order to avoid any confounding changes in gene expression, which can affect R-loop levels^20,41^ and AAV-HR integration^12^ (Fig. 4e). Firstly, we transduced HEPA 1-6 cells with either AAV- HR_neg_-mAlb/Fluc or AAV-HR-mAlb/Fluc. Remarkably, unlike the AAV-HR-mAlb/Fluc, AAV- HR_neg_-mAlb/Fluc treated cells did not exhibit any detectable luciferase activity or mRNA production despite the presence of AAV genomic DNA into cells (Fig. 4f, Extended Data Fig. 4c). As expected, topotecan treatment did not alter AAV-HR_neg_-mAlb/Fluc expression in HEPA 1-6 (Extended Data Fig. 4d).

Finally, we treated mice with AAV-HR_neg_-mAlb/hFIX and AAV-HR-mAlb/hFIX. Consistent with our data from HEPA 1-6 cells, AAV-HR_neg_-mAlb/hFIX did not integrate into murine genome, as assessed by the lack of detectable circulating hFIX, compared to a time-dependent increase detected in animals treated with AAV-HR-mAlb/hFIX (Fig. 4g). Moreover, analysis of AAV vector copy number in liver demonstrated that there were no differences in AAV transduction efficiencies between AAV-HR_neg_-mAlb/hFIX and AAV-HR-mAlb/hFIX vectors (Fig. 4h).

These results indicate that AAV-mediated genome integration in mice was promoted by R-loop formation at the 3’ end of the *Alb* locus.

### AAVs may preferentially integrate in transcriptionally active loci with high levels of R-loops

In recent years, more groups have been sequencing the entire liver genome after systemic AAV-mediated gene editing to assess potential off-target vector integrations. Chandler et al. found that AAV integration in the murine-specific *Rian* locus induced hepatocarcinoma in mice. However, the AAV vector was also found integrated in highly transcribed loci such as *Alb* and *Tfr*^42^. Moreover, a recent study aiming to target the *TCR* locus with AAV, found integration to be mostly within the *Alb* locus upon systemic vector delivery^43^. On the other hand, researchers aiming to target the *Alb* locus by CRISPR/Cas9-mediated HR discovered secondary off-target genes in *Tfr* and *Afp*^44^. More recently, De Giorgi et al., used CRISPR/Cas9-mediated HR to perform genome editing in the hepatic *Apoa1* locus and found integrated AAV vectors again in *Alb*, *Afp*, and *Errfi1* loci^45^. Interestingly, the genes that are prone to off-target AAV integration have high R-loop levels (Fig.5a), although, the most frequent “off-target” gene in the liver is *Alb* which is also the most highly transcribed locus (Fig. 5b). Moreover, despite finding a positive correlation between R-loops formation and AAV integration, additional studies are needed on whether this mechanism influences the NHEJ DNA repair (known to affect off-target/random AAV insertions) or the HR pathway which underlie the AAV-HR gene editing.

**Figure 5:**
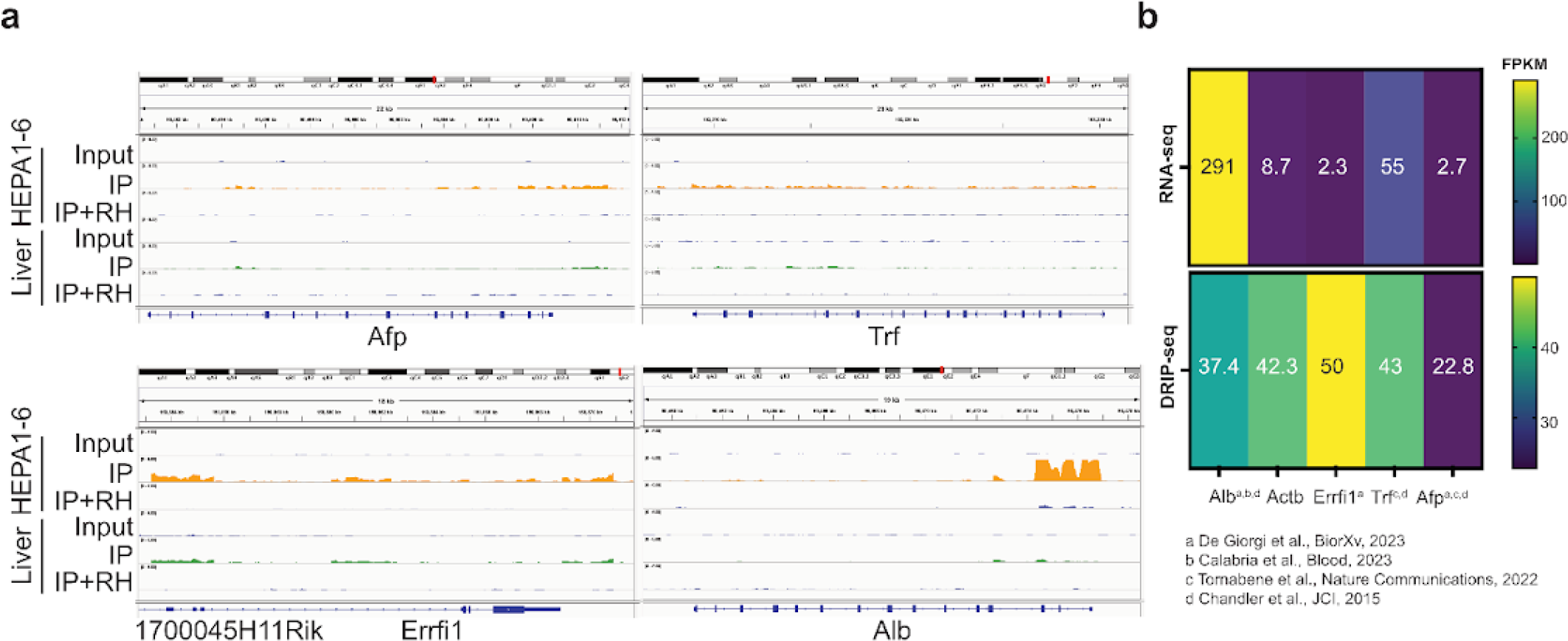
Correlation between off-target genes and R-loops a. IGV visualization of previously reported genes where AAV vectors were found integrated upon in vivo liver gene transfer. **b.** Heatmap showing DRIP-seq and RNA-seq values of genes represented in **a.**

Thus, while transcription may be the main determinant of AAV genome integration, co-transcriptional R-loop formation may also play a significant role in influencing the sites of vector integration in the host genome.

## Discussion

This study reveals an unanticipated role of genomic R-loops in AAV-mediated genome editing. Specifically, genome-wide R-loop mapping in hepatoma HEPA1-6 cells and mouse liver allowed us to identify genomic regions with a combination of high R-loop and transcription levels that are prone to AAV vector integration. Differences in transcription and R-loop formation between the two cell types we studied underscores the importance of being able to map R-loops genome-wide in primary tissue (*in vivo* DRIP-seq), although immortalized cell lines provide a useful system for studying the effects of R-loops on AAV-HR.

In this context, our data in primary murine liver confirm previous studies in immortalized cells, showing that DRIP peaks are mostly present in transcribed genes at both the 5’ (transcription starting site) and 3’ (transcription termination site) end regions of genes (Fig. 3 and Extended Data Fig. 2). Of note, in the *Alb* locus, we observed a robust RNA-DNA hybrid signal at the 3’ end with very low or no detectable signal towards the 5’ end of the gene. In our experiments, we demonstrated that the high levels of RNA-DNA hybrids at the 3’ end of *Alb* favored nuclease-free genome integration of an AAV with arms homologous to the *Alb* transcription termination site. Importantly, the genetic and pharmacological inhibition of factors which resolve or suppress RNA-DNA hybrids led to a significant increase in AAV-HR at the *Alb* locus. Conversely, overexpression of RNaseH1, an enzyme that specifically degrades RNA-DNA hybrids, reduced AAV-HR. Remarkably, our attempt to perform AAV-HR at an *Alb* region devoid of RNA-DNA hybrids failed to show any detectable genome editing events both *in vitro* and *in vivo*. Given that *Alb* is one of the most highly transcribed genes, it would be interesting to investigate whether the preferential accumulation of R-loops at the gene terminus could be used as a specific molecular signature for in vivo genome editing approaches. It is possible that additional factors may be important, for example certain nucleotide sequences or DNA secondary structures that contribute to formation of highly stable R-loops^23,27^.

We propose a general mechanism of AAV vector integration in liver-directed gene transfer (Fig. 6). High levels of gene expression and consequent co-transcriptional accumulation of RNA-DNA hybrids might trigger the AAV integration through annealing of the single stranded AAV genome and the displaced DNA strand in R-loops. Vector inverted terminal repeats, known for their recombinogenic properties^10^, might also facilitate homologous recombination with R-loops present in the host genome and facilitate AAV integration. Alternatively, regions that are highly transcribed and R-loop prone can be sources of endogenous DNA replication stress and DNA damage, including some gene terminators which are sites of potential conflicts between transcription and replication machinery^34^. Such spontaneous DNA breaks may promote AAV integration by transcription-associated recombination, a process that efficiently repairs DNA lesions at transcriptionally active loci. R-loops have been reported to stimulate homologous recombination associated with DNA breaks at transcribed regions^25,46^. Additionally, chromatin state associated with DNA transcription has been suggested as determinant of AAV-HR and vector genomic integration^11^.

**Figure 6:**
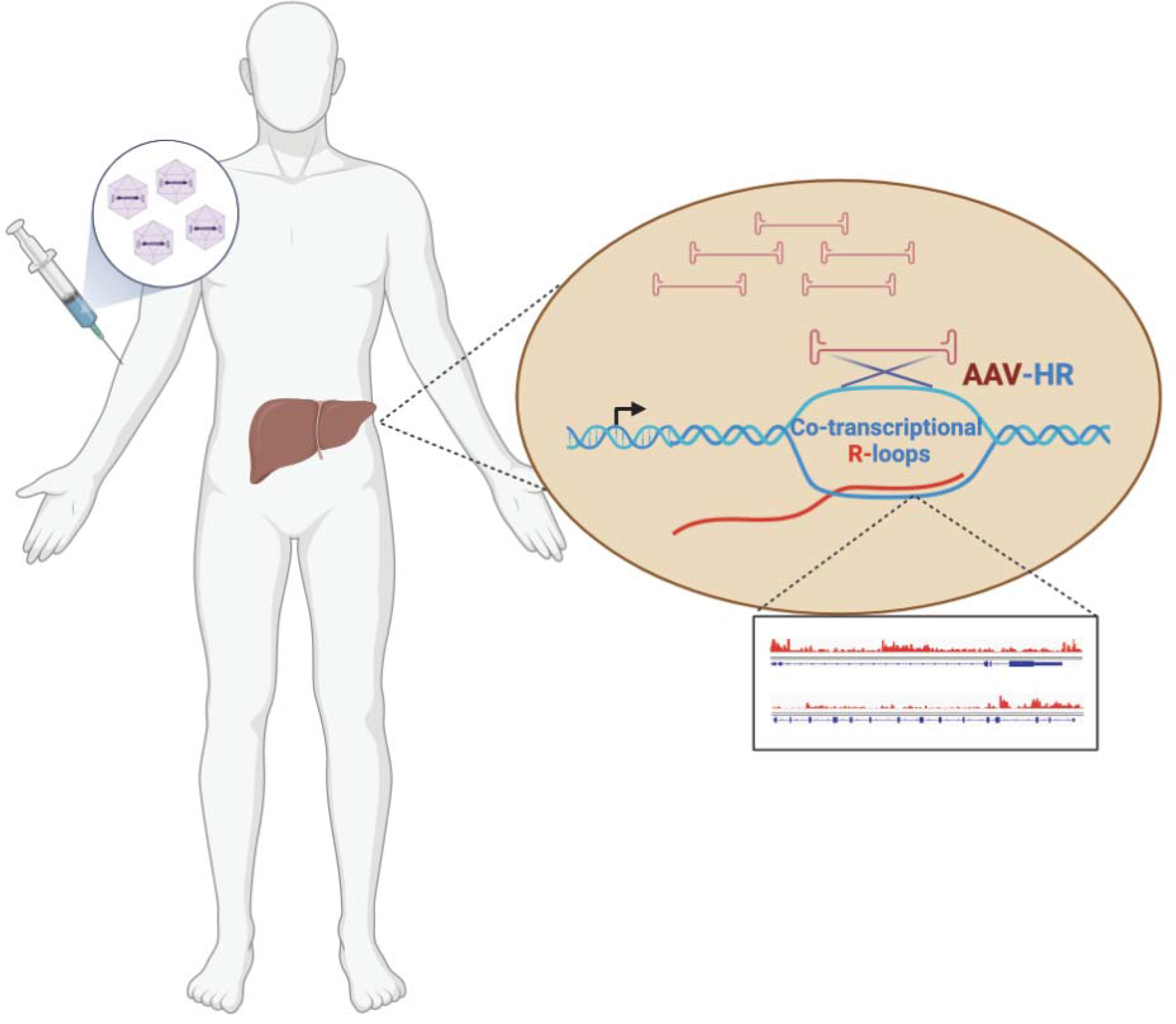
Mechanism of AAV-mediated integration at R-loop-forming loci. Systemic treatment with AAV for liver-mediated gene therapy. The single-stranded AAV genome is released into the nucleus of hepatocytes, undergoing homologous recombination (HR) with the single-stranded genomic DNA displaced by R-loop formation in transcriptionally active regions.

AAVs have been described as non-integrative vectors by forming episomal concatemers in the cell nucleus to establish durable expression of their cargo genes^3^. However, this view has been challenged with reports showing low frequencies of random AAV vector genome integration upon *in vivo* liver gene transfer in different species^8,47,48^. Moreover, independent studies described how AAVs have been found within multiple genes throughout the murine liver genome^42–45^. Interestingly, in our DRIP-seq experiments in mouse liver we found out that most of these ’off-target’ integrations genes exhibit high levels of R-loops (Fig. 5) . Of note, the *Alb* locus was the most common ’off-target’ genome region where the vector was found integrated in all the studies taken into consideration^42–45^. Thus, we speculate that a combination of high transcription (*Alb* is the highest transcribed gene in liver) and R-loop levels might promote AAV vector genome integration, without ruling out the possibility that this combination of events could also lead to increased DNA damage which in turn could also favor AAV integration^49^. Moreover, R-loops are also known to affect the major DNA repair pathways^50,51^, therefore whether R-loop formation influences both HR-mediated AAV insertion, and NHEJ-driven random integration events require further investigation.

To our knowledge, this represents the first DRIP-seq study using intact mammalian tissue. Previous studies have performed DRIP-seq on tissue samples, but these commonly use fresh tissue and extract genomic DNA from single cells^52^. Our modified approach allows for processing of frozen tissue that can be easily stored. This DRIP-seq approach may facilitate studies into the roles of R-loops in disease by being able to map genome-wide changes in RNA-DNA hybrids in animal models of disease, rather than being largely inferred from cell lines^18^. We envision that comparing DRIP-seq datasets from primary and transformed tissues *in vivo* could reveal additional roles of RNA-DNA hybrids in the initiation of cancer^53^. Moreover, DRIP-seq data obtained in brains from murine models of neurological disorders, in which R-loops are reported to have a role in the disease pathogenesis could improve the in vivo understanding of these disorders^54^. Mapping R-loops in different stages of development in primary tissues may also address the roles of RNA-DNA hybrids in aging^55^. Finally, sequencing RNA-DNA hybrids in vivo may improve our understanding of the role of hybrids in inflammation and autoimmune disorders^23,56,57^.

The gene editing systems based on CRISPR components are all dependent on RNA-DNA hybrid formation between the guide RNAs and genomic DNA, which is required for activation of the Cas9 enzyme^58^. Intriguingly, a recent study has reported that negatively supercoiled DNA increases genome-wide CRISPR/Cas9 off-target cleavage^59^, a property of DNA also known to strongly promote R-loop formation^60^. Thus, *in vivo* DRIP-seq might be important for evaluating the efficacy of Cas9-mediated genome editing, in addition to AAV approaches. Collectively, our study identifies R-loop formation as an important genomic factor that influences AAV-HR, improving our understanding of AAV integration. Being able to better predict regions prone to insertional mutagenesis when employing viral vectors for therapeutic approaches could ultimately help improve the safety and efficacy of genome editing.

## Material and methods

### Plasmid construction

The pAAV plasmids generated in this study were produced by Gibson assembly using NEBuilder® HiFi DNA Assembly Master Mix (NEB, E2621S). The AAV-HR for in vitro studies in HEPA1-6 cells was obtained by replacing the hFIX gene^5^ with the firefly luciferase (Fluc) cDNA (devoid of the starting codon (ATG)) contained in the plasmid we previously produced in our lab and also available from Addgene (plasmid #83281). The AAV-HR targeting the negative region of R-loops (AAV-HR_neg_) of the albumin locus was constructed by topo cloning (Thermo Fisher, 451245) while the homology arms were PCR amplified from gDNA extracted from murine liver (DNA plasmid sequences can be found in the supplementary material section).

The AAV-HR vector used for in vivo studies has been previously characterized^5^. The *ppyCAG_RNaseH1_WT* employed for the transient overexpression of RNAseH1 in vitro on HEPA1-6 cells was a gift from Xiang-Dong Fu (Addgene plasmid #111906).

### AAV vector production

Transfection for AAV vectors production was performed as previously reported^61^. For in vitro experiments the AAV vectors were purified using AAVpro® Purification Kit (Takara, 6666) following manufacturer instructions. For the studies in vivo the AAV were purified by CsCl gradient as already described^62^.

### In vitro studies

HEPA1-6 cells were grown in media supplemented with 10% fetal bovine serum (FBS), 2 mM L-glutamine, 100 IU/ml penicillin/streptomycin, in a humidified incubator at 37°C with 5% CO2.

HEPA1-6 transduction studies were performed by adding to the cells the AAV-HR (AAVDJ- mAlb/Fluc) at a final MOI of 50,000vg, unless stated otherwise in the figure legends.

After 48h, cell lysis and luciferase assay were performed using the Promega Luciferase 1000 Assay System (Promega, E4550) following manufacturer instructions.

For the knockdown experiments, all siRNAs were acquired from Dharmacon and resuspended in nuclease-free water to a concentration of 20 μM prior use:

-siRNA-Fancm (SMART pool: ON-TARGETplus Mouse Fancm siRNA (L-058186-01-0005))

-siRNA-Srsf1 (SMART pool: ON-TARGETplus Mouse Srsf1 siRNA (L-040886-01-0005))

-siRNA-Top1 (SMARTpool: ON-TARGETplus Mouse Top1 siRNA (L-047567-01-0005))

-siRNA-Top3b (SMARTpool: ON-TARGETplus Mouse Top3b siRNA (L-062791-01-0005))

-siRNA-Sin3a (SMARTpool: ON-TARGETplus Mouse Sin3a siRNA (L-044653-01-0005))

-siRNA-Rnaseh1 (SMARTpool: ON-TARGETplus Mouse Rnaseh1 siRNA (L-043445-02- 0005))

-siRNA-scramble (ON-TARGETplus non-targeting siRNA #1 (D-001810-01-50))

The day before transfections, HEPA1-6 cells were seeded. The next day, siRNAs were transfected (at a final concentration of 100nM) with the Dharmafect 4 solution (Dharmacon, T- 2004-02) following manufacturer instructions. After 48h cells were transduced with the appropriate AAV-HR vector. Luciferase activity was assessed 48h after AAV transduction.

In vitro transient overexpression of human RNaseH1 was achieved by transfecting the *ppyCAG-RNaseH1* plasmid. The day before the plasmid transfection, HEPA1-6 cells were seeded and on the next day, the plasmid was transfected using Lipofectamine 3000 (Thermo Fisher, L3000015) following manufacturer instructions. After 48h the cells were transduced with the AAV-HR vector. Luciferase activity was assessed 48h after AAV transduction.

In vitro drugs experiments were performed by adding Topotecan (Fisher Scientific, AC467180010) to the HEPA1-6 at the final concentration of 1μM. After 1 hour the cells were washed one time with DPBS and transduced with AAV-HR vectors. In vitro experiment using Fludarabine was conducted as previously described^39^ by incubating the cells for 16 h with 100 nM of drug before adding the AAV-HR vector. Luciferase activity was assessed 48h after AAV transduction.

For western blot analysis, HEPA1-6 were lysed in RIPA buffer and loaded on a 4-15% gradient polyacrylamide gel (Thermo Fisher, NP0335). Protein transfer was carried out using the iBlot system (Thermo Fisher, IB23002). The nitrocellulose membrane was blocked with Odyssey buffer (Fisher, NC0730870) and incubated with the anti-RNaseH1 (GeneTex, GTX117624) and the anti-tubulin (Sigma, T9026) antibody both diluted to 1:1000 in odyssey buffer. The membrane was washed and incubated with an anti-rabbit secondary antibody (Fisher, NC9401842) or anti-mouse secondary antibody (Fisher, NC0809365) diluted 1:5000 and visualized by Odyssey imaging system (Li-Cor Biosciences).

mRNA analyses were performed by extracting total mRNA from HEPA1-6 cells using RNeasy mini kit (Qiagen, 74104) following manufacturer instructions. mRNA was converted to cDNA using Maxima First Strand cDNA Synthesis Kit for RT-qPCR (Thermo Fisher, K1671).

For vector copy number analysis, total gDNA was extracted using a QIAamp DNA Mini kit (Qiagen, 51304) according to the manufacturer instructions. Standard curves were generated using serially diluted linearized plasmid and used for vector copy number quantification.

### qPCR

mRNA, vector copy number analysis, AAV titration, and DRIP-qPCR were carried out by qPCR in the CFSX384 instrument (Biorad) using Brilliant III Ultra-Fast SYBR QPCR MM (Agilent Technologies, 600882). Primers used for qPCR experiments are listed in Supplementary table 2.

### In vivo studies

Mouse experiments were conducted and approved by the Administrative Panel on Laboratory Animal Care of Stanford University. 6–8-week-old C57/Bl6 mice (JAX, 000664) were kept in the animal facility with a normal night/day cycle and regular chow *ad libitum*. AAV-HR vectors were intravenously injected at the dose of 3x10^11^ vg/mouse. Standard ELISA for the detection of circulating hFIX was performed. Briefly, 96-well plates (Thermo Fisher Scientific, 442404) were coated with 1 μg/mL of anti-hFIX antibody (Sigma, F2645) in coating buffer (10 mM phosphate buffer, pH 7.4; Sigma, C3041). Human FIX standard curve was prepared using commercial purified hFIX (AbCam, ab62544). The plates were incubated overnight at 4°C, washed (wash buffer, PBS/Tween 20 0,05%), blocked at room temperature in blocking buffer (PBS/BSA 6%) for one hour, and incubated with the serum samples for 2h at 37°C. After washing, plates were incubated with HRP-conjugated goat anti-hFIX (diluted 1:5,000; Fisher, NC1762057) for one hour at 37°C. The plates were then washed, incubated with the substrate o-phenylenediamine dihydrochloride, OPD (Sigma, 34006), for 5-10 minutes at room temperature; the reaction was stopped by adding 1 volume of H_2_SO_4_ (2.5N) and the plates were read at a wavelength of 492 nm using TECAN reader.

Total gDNA for AAV vector copy number analysis was extracted using Puregene Tissue Kit (Qiagen, 158063) according to the manufacturer instructions. Standard curves were generated using serially diluted linearized plasmid and used for vector copy number quantification.

Topotecan (SelleckChem, S1231) was resuspended in saline buffer (NaCl 0.9%). The mice were treated by intraperitoneal injection. The day of tissues collection the animals were deeply anesthetized and then transcardially perfused with PBS. Snap-frozen liver or brain were homogenized in two RINO 1.5-ml Screw-Cap tubes with 850Lμl per tube of PBS using Bullet Blender (Next Advance; Liver: 5’ at speed 10, Brain: 3’ at speed 8) at 4°C. The homogenate from the two RINO tubes was then pooled into a sterile 2mL tube. For DRIP protocol, 250μL (500μL for brain) of liver were washed three times with PBS by centrifugation (at 4°C 500g x 5’) and finally resuspended in 1.6mL of TE buffer to start the genomic DNA extraction for DRIP experiments^26^.

### DNA/RNA immunoprecipitation (DRIP)

DRIP-qPCR experiments were performed as previously described^26^. Briefly, genomic DNA was extracted from cells using proteinase K digestion followed by phenol:chloroform extraction. Genomic DNA was then digested with a cocktail of restriction enzymes (Bsp1407I, SspI, XbaI, EcoRI, HindIII). 8 μg of genomic DNA was used for each immunoprecipitation using the S9.6 antibody (Sigma, MABE1095). The immunoprecipitated DNA was then used for DRIP-qPCR analyses.

For the DRIP-seq experiments we used the same samples as for DRIP-qPCR, but the DNA sequencing libraries were prepared as previously described^20,27^.

Briefly, prior to library preparation DNA was sonicated in Covaris microtubes (Fisher, 520045) to a peak fragment size of 300 bp, performed on a Covaris machine S220 using the following parameters: Intensity: 4, C/B: 200. DF: 10%, Time: 30 seconds.

NEBNext® End Repair Module (NEB, E6050S) was used to repair the sonicated DNA and successively 3’ A-tails were added by Klenow Fragment (3’→5’ exo-) (NEB, M0212S) and dATPs (NEB, N0440S). Next, adapters were ligated to the repaired DNA using the 1S Plus Set A Indexing Kit (Swift Biosciences), and DNA was then amplified by PCR (PE PCR Primer 1.0: 5’AATGATACGGCGACCACCGAGATCTACACTCTTTCCCTACACGACGCTCTTCCGAT CT 3’; PCR Primer 2.0: 5’ CAAGCAGAAGACGGCATACGA 3). For input libraries, 4 PCR cycles were used, while for IP libraries, 10 PCR cycles were needed. Following PCR amplification, DNA fragments between 200 and 600 bp were size selected using AMPure XP beads (Beckman Coulter, A63880). Library DNA was analyzed on a Bioanalyzer DNA HS (Agilent, Santa Clara, CA, USA) then pooled in equimolar ratios and sequenced on a HiSeq 4000 (Illumina, San Diego, CA, USA) at the Stanford Genome Sequencing Service Center, using 2x75bp sequencing.

### RNA-seq analyses

RNA-seq data analyses were performed using the data collected by the NCBI repository website. Raw FASTQ files for HEPA1-6 cell lines: <u>SRX14425594</u>, <u>SRX14425595</u>, <u>SRX14425599</u>.

Raw FASTQ files for murine livers: SRX20088936, SRX20088937, SRX20088938.

The quality of the FASTQ files was assessed using FastQC/ v0.11.8 (https://www.bioinformatics.babraham.ac.uk/projects/fastqc/), checking for parameters such as the presence of overrepresented sequences, adapter contamination and per-sequence quality scores. Specific and universal adapter sequences were removed from each FASTQ files using Cutadapt v 1.18^63^. Once adapter sequences were trimmed, they were aligned to the Genome Reference Consortium Mouse Build 38 (GRCm38)^64^ using the STAR aligner (STAR/2.7.10b)^65^ with Gencode VM25 basic annotation as the reference gene annotation. Aligned reads were sorted using SAMtools/1.16.1^66^. Quantification of the reads was performed using the Subread featureCounts package (Subread/2.0.6)^67^, retaining reads greater than 0 for analysis. Differential gene expression analysis was conducted using Deseq2/3.18^68^, identifying differentially regulated genes with an absolute log fold change of 0.5 and a p-adjusted value of 0.01. Up and down- regulated genes were calculated using absolute log-fold changes of +0.5 and -0.5, respectively. From the raw counts generated by featureCounts, FPKM values for each gene were calculated along with gene length, chromosomal coordinates, and gene orientation.

### GO and KEGG enrichment of DEG using Cluster profiler

GO and KEGG enrichment of differentially expressed genes (DEG) across all murine liver and HEPA1-6 samples were performed using the Cluster Profiler package in R^69^. Enrichment analysis aimed to analyze specific enriched clusters in gene sets rather than individual genes, providing insight into specific GEO and KEGG terms. KEGG enrichment was conducted for individual gene sets, and specific enrichment scores were computed. Common GEO and KEGG terms across all libraries were identified, with the number of permutations set to 10,000, and minimum and maximum gene sets set to 3 and 800, respectively. The adjusted p-value cutoff was 0.05.

### Correlation analysis of RNA-seq data with DRIP-seq data

Raw FPKM values have been computed for both HEPA1-6 and liver cell data across three replicates for the RNA-seq data. Similar computations have been made for HEPA1-6 and liver cells for DRIP-seq data, and Pearson correlation values have been calculated for both RNA-seq data and DRIP-seq data. BEDTools coverage was used to calculate DRIP-seq read coverage over mouse genes (Gencode VM25 basic). FPKM DRIP-seq and RNA-seq values over the same genes were correlated in hexbin plots, showing gene densities. Correlations with R-squared values were computed. Genes intersecting an R-loop peak were called R-loop positive (+) and those without an R-loop peak were called R-loop negative (-).

### DRIP seq analysis

Cutadapt was used to remove adapter sequences, and trimmed reads were aligned to the murine genome reference mm10 using bowtie2 in end-to-end alignment mode and with unaligned, mixed and discordant reads excluded. Duplicate reads were removed with Picard. BEDTools was used to convert these SAM files into BEDPE format. Genome browser tracks were produced with the BEDTools genomecov utility, normalized to reads per million mapped, and visualized using IGV.

### Peak calling

Deduplicated bam files from biological replicates were merged using samtools and then peaks were called between these merged input and IP files using MACS2 with default broad peak settings. Peak strand annotations were assigned by intersecting peaks with GencodeVM25 gene annotations extended up- and downstream by 3 kb.

### Metaplots

Metaplots around genes, transcription start sites, transcription end sites and DRIP peaks were produced from genome browser tracks with deepTools. Tracks for GC and AT-skew were produced using the BEDTools *nuc* tool with further processing from unix text processing tools. To calculate density of non-B DNA structures around peak sites, the locations of predicted secondary structures turned into a track using the bedtools genomecov utility (where the track was 1 to indicate the presence of a motif, or 0 to indicate the absence), and these values were used to produce metaplots with deepTools. GC and AT content within peaks was calculated using BEDTools nuc. Analysis of genome compartments overlapping DRIP-seq peaks was performed using the *cis*-regulatory element annotation system. Data processing for all genomic plots was performed using Python, NumPy and Pandas. Data were visualized using the Python packages Matplotlib and Seaborn.

### Statistics

GraphPad Prism and Python SciPy were used for statistical analysis. Data were analyzed by parametric tests, α =0.05 (one-way and two-way ANOVA with Tukey’s or Sidak’s post hoc correction). Nonparametric tests were performed when only two groups were compared (unpaired t-test). The statistical analysis performed for each data set is indicated in the figure legends. The p-value is shown for each data set present in the graphs. Bars in graphs represent standard deviation for each group.

## Data availability

DRIP-seq data generated in this study are publicly available on the NCBI repository database under the GEO accession number GSE261759.

## Code availability

https://github.com/madziacrossley/mouse_drip-seq_liver_hepa

## Acknowledgements

We would like to thank Xuhuai Ji and John Collier at the Stanford Genomics Facility for their help with the DNA sequencing. This work was supported by grants from the National Institutes of Health, R01-HL064272 (M.A.K), S10OD018220 and 1S10OD021763 to the Stanford Functional Genomics Facility, GM119334 to K.A.C., T32GM136631 and the NSF DGE- 2146755 to J.L.G, and the Glenn Foundation for Medical Research to K.A.C. KAC is an ACS research professor.

## Author contributions

F.P. and M.A.K. conceived and directed the study. F.P. performed all the experiments, analyzed the data, and helped with the in vivo work. M.P.C. generated protocols for library preparation and analyzed the DRIP-seq and RNA-seq data. A.G. and J.L.G. helped analyze RNA-seq data.

F.P helped F.Z. with the in vivo work (injection, blood and tissue collection, and mice perfusion). K.P. provided the AAV-HR vector used for the in vivo experiment in Figure 4c.

K.A.C. and M.P.C. shared reagents and provided critical insights on R-loop biology. F.P.,

M.P.C. and M.A.K. contributed to the interpretation of results and the significance of the work.

F.P. and M.P.C. wrote the manuscript.

---

**Extended Data Figure 1:**
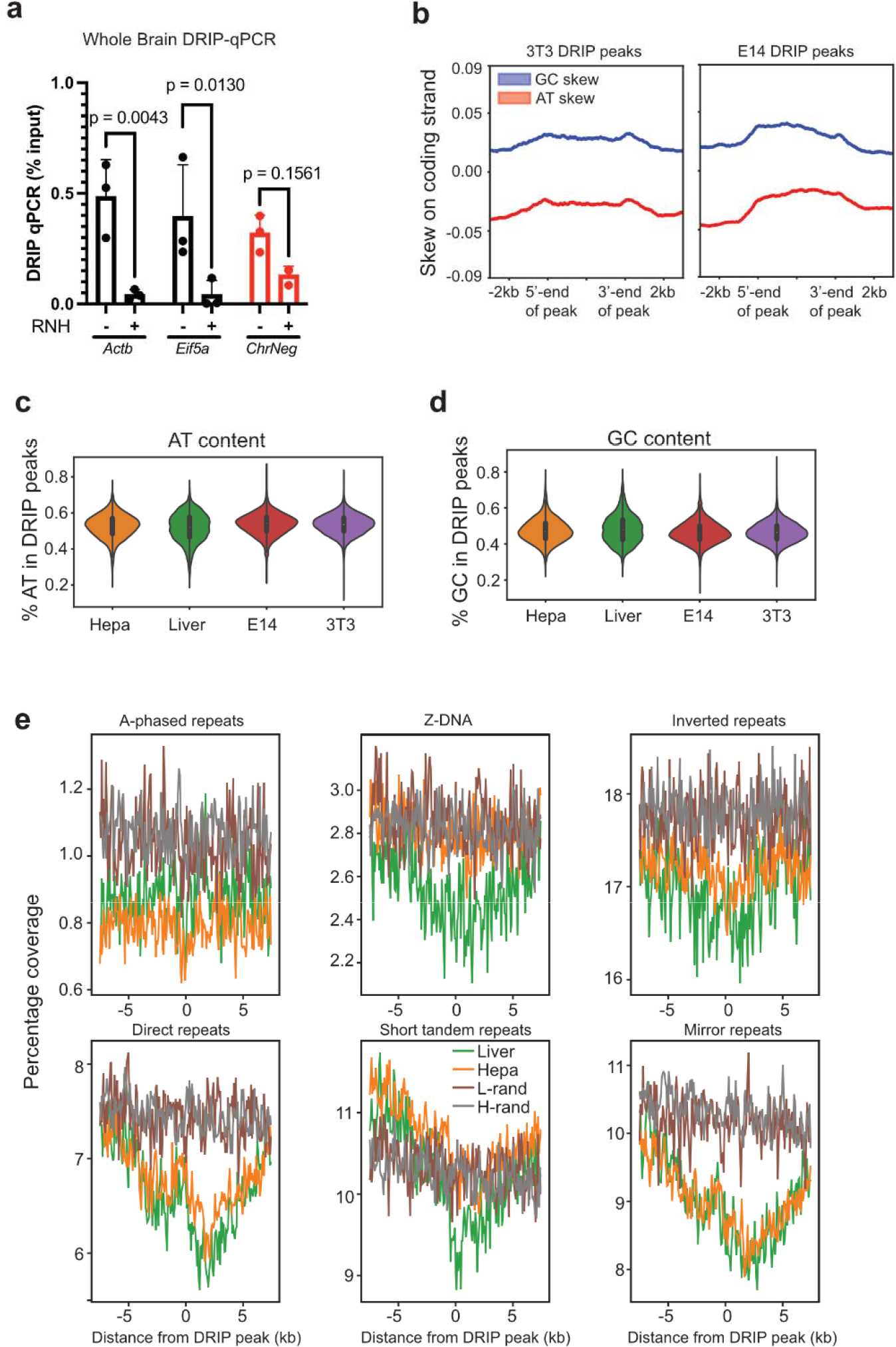
DRIP-seq analysis. a. DRIP-qPCR analysis in the whole mouse brain (n=3). **b.** GC (blue) and AT (red) skew across coding strand of DRIP peaks, including 2kb flanking 5’- and 3’-ends in 3T3 and E24 murine cells. Bands represent 95% CI of mean skew signal **c.** GC content percentage of DRIP peaks in HEPA1-6, liver, 3T3, and E14 cells. **d.** As in **c**, but for AT content. **e.** Aggregate plots around the center of DRIP peaks showing non-B DNA motif density as percent coverage Statistical analysis: **a.** Two-way ANOVA with Sidak’s post hoc analysis. Error bars represent standard deviation of the mean.

**Extended Data Figure 2:**
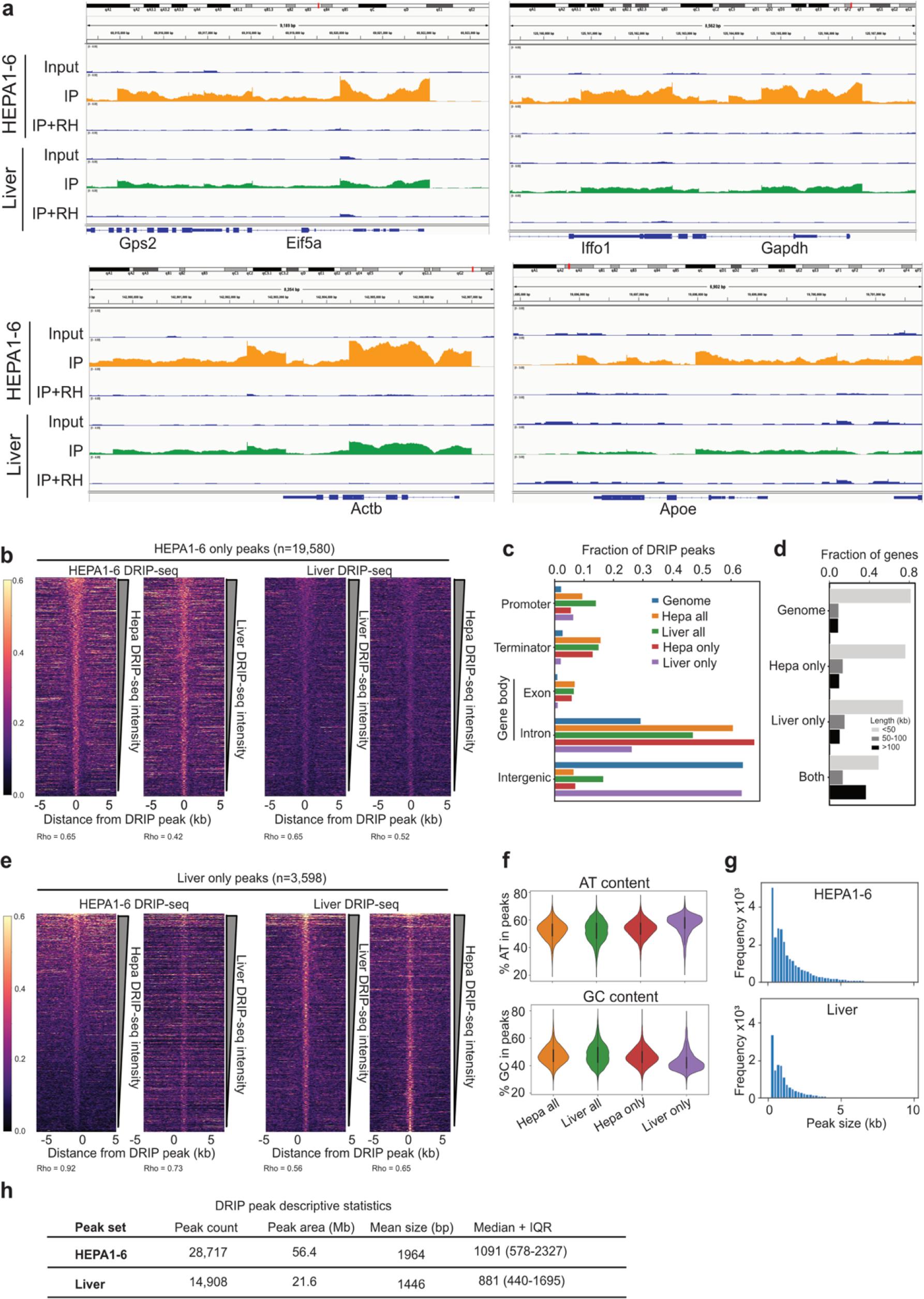
DRIP seq comparison HEPA1-6 vs. liver a. Genome browser views of DRIP-seq tracks of representative R-loop forming genes in HEPA1-6 and liver tissue. Tracks from S9.6 immunoprecipitation (IP), Input and IP with RNaseH treatment (IP+RH) are shown. **b.** Heatmaps showing the DRIP-seq signal around the center of peaks called uniquely in HEPA1-6 Heatmaps are ranked by DRIP-seq signal strength within 3kb of the peak center. Correlation coefficients (Spearman’s Rho) are shown **c.** Fractions of DRIP peaks from HEPA1-6 or liver overlapping murine genomic features. d. Lengths of genes overlapping DRIP peaks. Gene lengths for liver only peaks are significantly different to other peaksets (P=4.5e-20, Chi-square test). **e.** As in b, but for DRIP-seq signal around the center of peaks called uniquely in liver tissue. **f.** AT (top) and GC (bottom) content of DRIP peaks in HEPA1-6 and liver. **g.** Histogram of peak sizes obtained from peak calling on DRIP versus input samples. **h.** Table containing descriptive statistics of DRIP peaks in HEPA1-6 and liver.

**Extended Data Figure 3:**
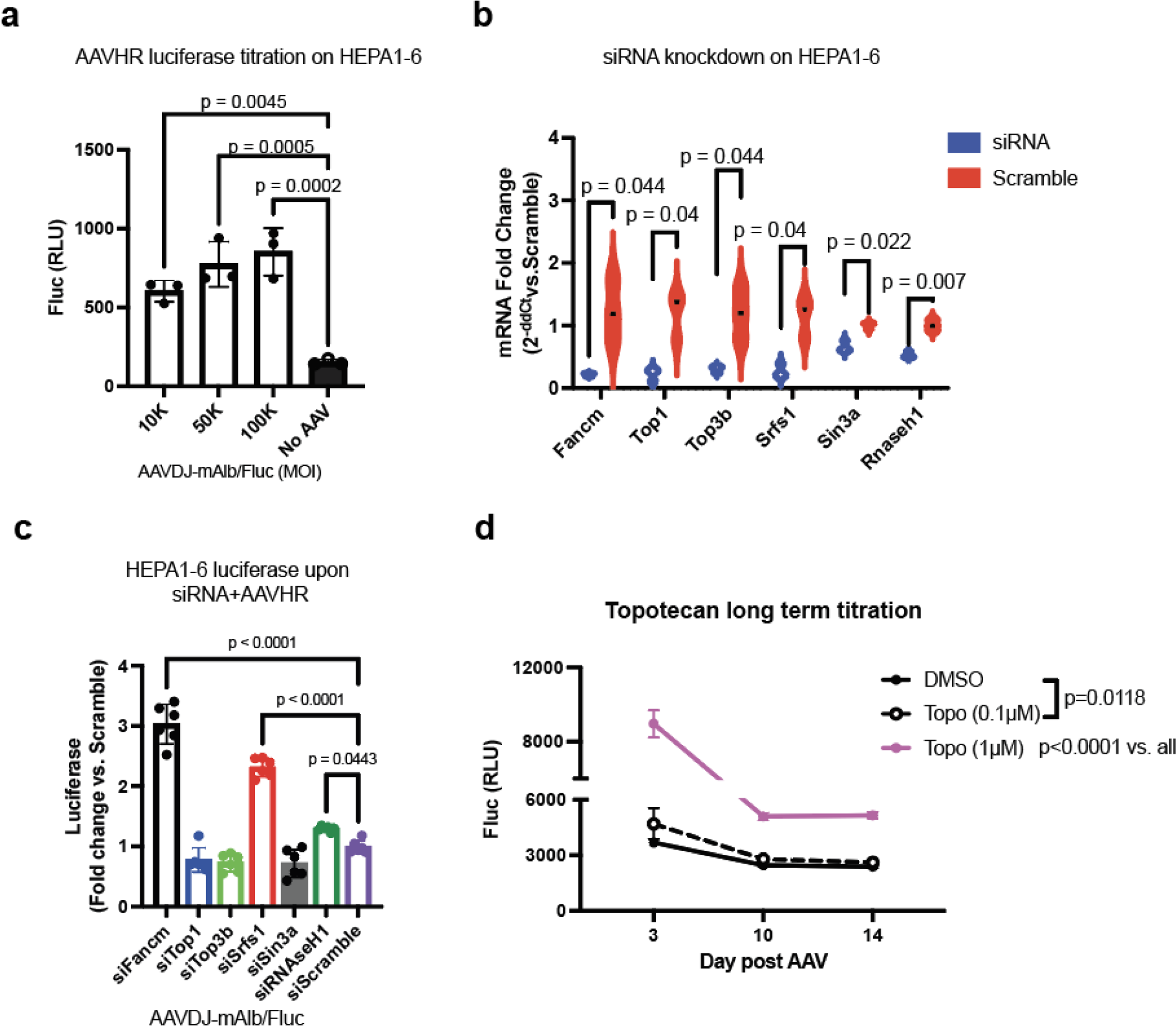
In vitro AAV-HR and R-loops a. Luciferase activity in HEPA1-6 upon AAVDJ-mAlb/Fluc transduction at different multiplicity of infection. Non transduced cells (No AAV) were used as negative control (n=3). **b.** mRNA levels upon transfection of siRNA against R-loops genes normalized against cells treated with Scramble siRNA (n=3). **c.** Luciferase activity in HEPA1-6 upon transfection of siRNAs against R-loops related proteins and AAVDJ-mAlb/Fluc transduction (n=6). Results were normalized to scramble-treated samples **d.** Time course of luciferase activity in HEPA1-6 upon treatment with either different doses of totopotecan or DMSO as negative control and transduced with AAVDJ- mAlb/Fluc (n=3). Statistical analysis: **a, c.** One-way ANOVA with **a.** Tukey’s and **c**. Sidak post hoc analysis**. b.** Multiple t-test using Holm-Sidak method. **d.** Two-way ANOVA with Sidak’s post hoc analysis. Error bars represent standard deviation of the mean.

**Extended Data Figure 4:**
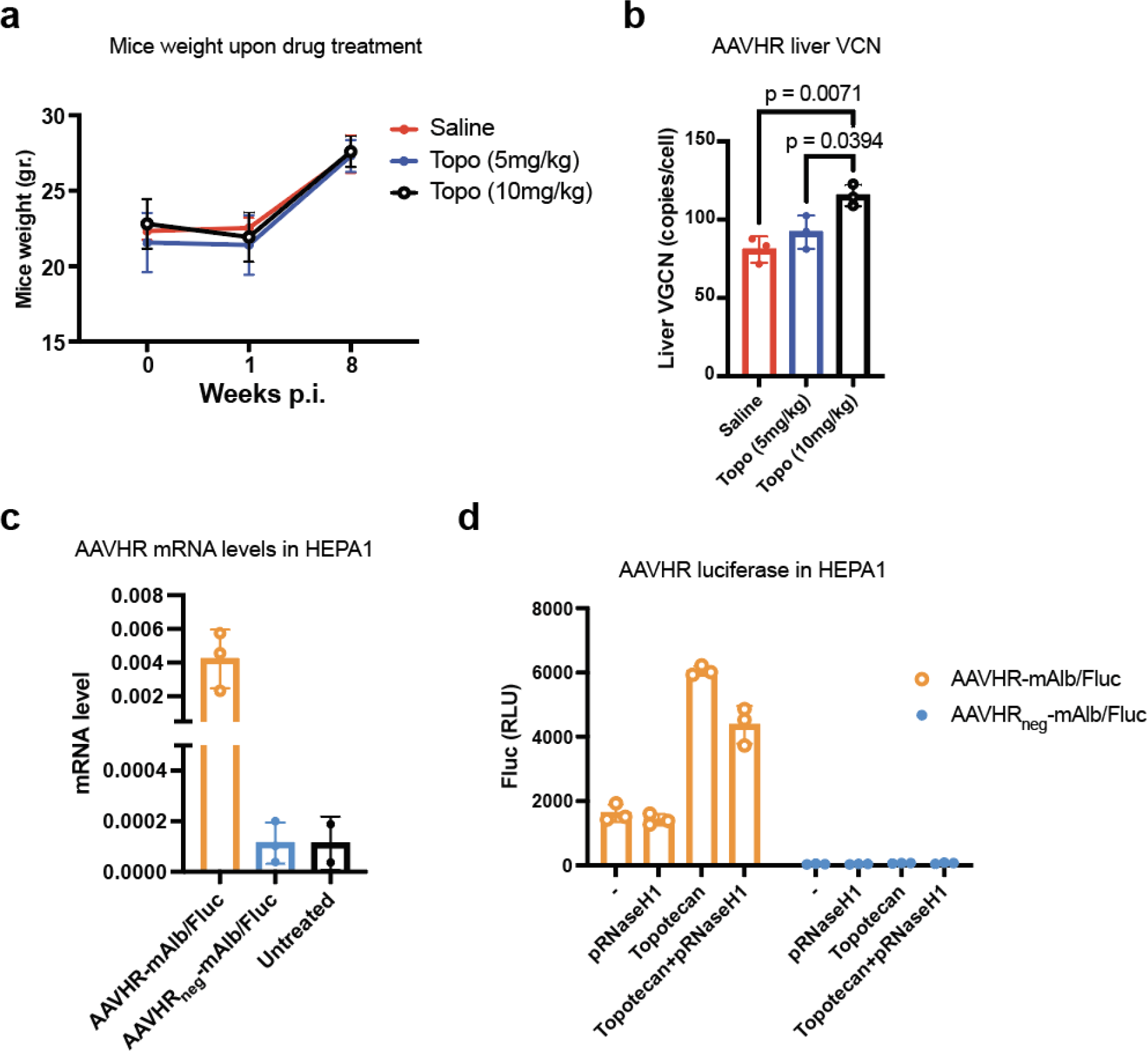
In vitro AAV-HR and R-loops a. Mice body weight upon treatment with different doses of topotecan and AAV-HR-mAlb/hFIX (n=3). **b.** AAV vector genome copy number in liver of mice treated with AAV-HR-mAlb/hFIX and different doses of topotecan (n=3). **c.** mRNA levels in HEPA1-6 transduced with AAV-HR- mAlb/Fluc and AAV-HR_neg_-mAlb/Fluc (n=3). **d.** Luciferase activity in HEPA1-6 upon the cells were treated with either totopotecan or pRNaseH1 plasmid or their combination, and AAV-HR- mAlb/Fluc or AAV-HR_neg_-mAlb/Fluc (n=3). Statistical analysis: **b.** One-way ANOVA with Tukey’s post hoc analysis. Error bars represent standard deviation of the mean.

## References

1. Li, C. & Samulski, R. J. Engineering adeno-associated virus vectors for gene therapy. Nature reviews. Genetics 21, 255–272 (2020).

2. Mendell, J. R. et al. Current Clinical Applications of In Vivo Gene Therapy with AAVs. Molecular Therapy 29, 1–25 (2020).

3. Nakai, H., Fuess, S., Storm, T. A., Meuse, L. A. & Kay, M. A. Free DNA ends are essential for concatemerization of synthetic double-stranded adeno-associated virus vector genomes transfected into mouse hepatocytes in vivo. Mol Ther 7, 112–121 (2003).

4. Wang, L., Wang, H., Bell, P., McMenamin, D. & Wilson, J. M. Hepatic gene transfer in neonatal mice by adeno-associated virus serotype 8 vector. Human gene therapy 23, 533–9 (2012).

5. Barzel, A. et al. Promoterless gene targeting without nucleases ameliorates haemophilia B in mice. Nature 517, 360–364 (2015).

6. Adikusuma, F. et al. Large deletions induced by Cas9 cleavage. Nature 560, E8–E9 (2018).

7. Miller, D. G., Petek, L. M. & Russell, D. W. Adeno-associated virus vectors integrate at chromosome breakage sites. Nature Genetics 36, 767–773 (2004).

8. Martins, K. M. et al. Prevalent and Disseminated Recombinant and Wild-Type Adeno- Associated Virus Integration in Macaques and Humans. Human Gene Therapy hum.2023.134 (2023) doi:10.1089/hum.2023.134.

9. Yan, Z., Zak, R., Zhang, Y. & Engelhardt, J. F. Inverted Terminal Repeat Sequences Are Important for Intermolecular Recombination and Circularization of Adeno-Associated Virus Genomes. J Virol 79, 364–379 (2005).

10. Hirsch, M. L. Adeno-associated virus inverted terminal repeats stimulate gene editing. Gene Therapy 22, 190–195 (2015).

11. Spector, L. P. et al. Evaluating the Genomic Parameters Governing rAAV-Mediated Homologous Recombination. Mol Ther 29, 1028–1046 (2021).

12. Nakai, H. et al. AAV serotype 2 vectors preferentially integrate into active genes in mice. Nat Genet 34, 297–302 (2003).

13. Greig, J. A. et al. Integrated vector genomes may contribute to long-term expression in primate liver after AAV administration. Nat Biotechnol 1–11 (2023) doi:10.1038/s41587-023-01974-7.

14. Fattah, F. J., Lichter, N. F., Fattah, K. R., Oh, S. & Hendrickson, E. A. Ku70, an essential gene, modulates the frequency of rAAV-mediated gene targeting in human somatic cells. Proceedings of the National Academy of Sciences of the United States of America 105, 8703–8 (2008).

15. Vasileva, A., Linden, R. M. & Jessberger, R. Homologous recombination is required for AAV-mediated gene targeting. Nucleic Acids Research 34, 3345–3360 (2006).

16. de Alencastro, G. et al. Improved Genome Editing through Inhibition of FANCM and Members of the BTR Dissolvase Complex. Molecular Therapy 29, 1016–1027 (2021).

17. Yang, S., Winstone, L., Mondal, S. & Wu, Y. Helicases in R-loop Formation and Resolution. J Biol Chem 105307 (2023) doi:10.1016/j.jbc.2023.105307.

18. Crossley, M. P., Bocek, M. & Cimprich, K. A. R-Loops as Cellular Regulators and Genomic Threats. Molecular Cell 73, 398–411 (2019).

19. Hamperl, S. & Cimprich, K. A. The contribution of co-transcriptional RNA:DNA hybrid structures to DNA damage and genome instability. DNA Repair (Amst) 19, 84–94 (2014).

20. Stork, C. T. et al. Co-transcriptional R-loops are the main cause of estrogen-induced DNA damage. eLife 5, e17548 (2016).

21. Hamperl, S., Bocek, M. J., Saldivar, J. C., Swigut, T. & Cimprich, K. A. Transcription- Replication Conflict Orientation Modulates R-Loop Levels and Activates Distinct DNA Damage Responses. Cell 170, 774–786.e19 (2017).

22. Sollier, J. et al. Transcription-Coupled Nucleotide Excision Repair Factors Promote R- Loop-Induced Genome Instability. Molecular Cell 56, 777–785 (2014).

23. Crossley, M. P. et al. R-loop-derived cytoplasmic RNA-DNA hybrids activate an immune response. Nature 613, 187–194 (2023).

24. Brickner, J. R., Garzon, J. L. & Cimprich, K. A. Walking a tightrope: The complex balancing act of R-loops in genome stability. Mol Cell 82, 2267–2297 (2022).

25. Ouyang, J. et al. RNA transcripts stimulate homologous recombination by forming DR- loops. Nature 594, 283–288 (2021).

26. Sanz, L. A. & Chédin, F. High-resolution, strand-specific R-loop mapping via S9.6-based DNA–RNA immunoprecipitation and high-throughput sequencing. Nature Protocols 14, 1734– 1755 (2019).

27. Crossley, M. P., Bocek, M. J., Hamperl, S., Swigut, T. & Cimprich, K. A. qDRIP: a method to quantitatively assess RNA–DNA hybrid formation genome-wide. Nucleic Acids Research 48, e84–e84 (2020).

28. Thongthip, S., Carlson, A., Crossley, M. P. & Schwer, B. Relationships between genome- wide R-loop distribution and classes of recurrent DNA breaks in neural stem/progenitor cells. Sci Rep 12, 13373 (2022).

29. Cer, R. Z. et al. Non-B DB v2.0: a database of predicted non-B DNA-forming motifs and its associated tools. Nucleic Acids Res 41, D94–D100 (2013).

30. Varshney, D., Spiegel, J., Zyner, K., Tannahill, D. & Balasubramanian, S. The regulation and functions of DNA and RNA G-quadruplexes. Nat Rev Mol Cell Biol 21, 459–474 (2020).

31. Dumelie, J. G. & Jaffrey, S. R. Defining the location of promoter-associated R-loops at near-nucleotide resolution using bisDRIP-seq. eLife 6, e28306 (2017).

32. Bader, A. S. & Bushell, M. DNA:RNA hybrids form at DNA double-strand breaks in transcriptionally active loci. Cell Death and Disease 11, (2020).

33. Chen, L. et al. R-ChIP Using Inactive RNase H Reveals Dynamic Coupling of R-loops with Transcriptional Pausing at Gene Promoters. Molecular Cell 68, 745–757.e5 (2017).

34. Promonet, A. et al. Topoisomerase 1 prevents replication stress at R-loop-enriched transcription termination sites. Nat Commun 11, 3940 (2020).

35. Parajuli, S. et al. Human ribonuclease H1 resolves R-loops and thereby enables progression of the DNA replication fork. Journal of Biological Chemistry 292, 15216–15224 (2017).

36. Salas-Armenteros, I. et al. Human THO-Sin3A interaction reveals new mechanisms to prevent R-loops that cause genome instability. EMBO J 36, 3532–3547 (2017).

37. Zhang, T. et al. Loss of TOP3B leads to increased R-loop formation and genome instability. Open Biology 9, (2019).

38. Pommier, Y. Topoisomerase I inhibitors: camptothecins and beyond. Nat Rev Cancer 6, 789–802 (2006).

39. Tsuji, S. et al. Fludarabine increases nuclease-free AAV- and CRISPR/Cas9-mediated homologous recombination in mice. Nat Biotechnol 40, 1285–1294 (2022).

40. Guichard, S. et al. Schedule-dependent activity of topotecan in OVCAR-3 ovarian carcinoma xenograft: pharmacokinetic and pharmacodynamic evaluation. Clin Cancer Res 7, 3222–3228 (2001).

41. Wahba, L., Costantino, L., Tan, F. J., Zimmer, A. & Koshland, D. S1-DRIP-seq identifies high expression and polyA tracts as major contributors to R-loop formation. Genes Dev. 30, 1327–1338 (2016).

42. Chandler, R. J. et al. Vector design influences hepatic genotoxicity after adeno-associated virus gene therapy. J Clin Invest 125, 870–880 (2015).

43. Calabria, A. et al. Intrathymic AAV delivery results in therapeutic site-specific integration at TCR loci in mice. Blood 141, 2316–2329 (2023).

44. Tornabene, P. et al. Therapeutic homology-independent targeted integration in retina and liver. Nat Commun 13, 1963 (2022).

45. Giorgi, M. D. et al. In vivo expansion of gene-targeted hepatocytes through transient inhibition of an essential gene. 2023.07.26.550728 Preprint at 10.1101/2023.07.26.550728 (2023).

46. Cohen, S. et al. Senataxin resolves RNA:DNA hybrids forming at DNA double-strand breaks to prevent translocations. Nat Commun 9, 533 (2018).

47. Nguyen, G. N. et al. A long-term study of AAV gene therapy in dogs with hemophilia A identifies clonal expansions of transduced liver cells. Nat Biotechnol 39, 47–55 (2021).

48. Sabatino, D. E. et al. Evaluating the state of the science for adeno-associated virus integration: An integrated perspective. Mol Ther 30, 2646–2663 (2022).

49. Guha, S. & Bhaumik, S. R. Transcription-coupled DNA double-strand break repair. DNA Repair (Amst) 109, 103211 (2022).

50. Petermann, E., Lan, L. & Zou, L. Sources, resolution and physiological relevance of R- loops and RNA–DNA hybrids. Nat Rev Mol Cell Biol 23, 521–540 (2022).

51. Audoynaud, C., Vagner, S. & Lambert, S. Non-homologous end-joining at challenged replication forks: an RNA connection? Trends in Genetics 37, 973–985 (2021).

52. Zhang, X. et al. Attenuation of RNA polymerase II pausing mitigates BRCA1-associated R-loop accumulation and tumorigenesis. Nat Commun 8, 15908 (2017).

53. Wells, J. P., White, J. & Stirling, P. C. R Loops and Their Composite Cancer Connections. Trends in Cancer 5, 619–631 (2019).

54. Perego, M. G. L., Taiana, M., Bresolin, N., Comi, G. P. & Corti, S. R-Loops in Motor Neuron Diseases. Mol Neurobiol 56, 2579–2589 (2019).

55. Schumacher, B., Pothof, J., Vijg, J. & Hoeijmakers, J. H. J. The central role of DNA damage in the ageing process. Nature 592, 695–703 (2021).

56. Giordano, A. M. S. et al. DNA damage contributes to neurotoxic inflammation in Aicardi-Goutières syndrome astrocytes. Journal of Experimental Medicine 219, e20211121 (2022).

57. Cristini, A. et al. RNase H2, mutated in AicardiLGoutières syndrome, resolves co- transcriptional R-loops to prevent DNA breaks and inflammation. Nat Commun 13, 2961 (2022).

58. Pacesa, M. et al. R-loop formation and conformational activation mechanisms of Cas9. Nature 609, 191–196 (2022).

59. Newton, M. D. et al. Negative DNA supercoiling induces genome-wide Cas9 off-target activity. Molecular Cell 83, 3533–3545.e5 (2023).

60. Chedin, F. & Benham, C. J. Emerging roles for R-loop structures in the management of topological stress. J Biol Chem 295, 4684–4695 (2020).

61. Grimm, D., Pandey, K., Nakai, H., Storm, T. A. & Kay, M. A. Liver transduction with recombinant adeno-associated virus is primarily restricted by capsid serotype not vector genotype. Journal of virology 80, 426–39 (2006).

62. Pekrun, K. et al. Using a barcoded AAV capsid library to select for clinically relevant gene therapy vectors. JCI Insight 4, e131610, 131610 (2019).

63. Martin, M. Cutadapt removes adapter sequences from high-throughput sequencing reads. EMBnet.journal 17, 10–12 (2011).

64. Mouse Genome Sequencing Consortium et al. Initial sequencing and comparative analysis of the mouse genome. Nature 420, 520–562 (2002).

65. Dobin, A. et al. STAR: ultrafast universal RNA-seq aligner. Bioinformatics 29, 15–21 (2013).

66. Li, H. et al. The Sequence Alignment/Map format and SAMtools. Bioinformatics 25, 2078–2079 (2009).

67. Putri, G. H., Anders, S., Pyl, P. T., Pimanda, J. E. & Zanini, F. Analysing high- throughput sequencing data in Python with HTSeq 2.0. Bioinformatics 38, 2943–2945 (2022).

68. Love, M. I., Huber, W. & Anders, S. Moderated estimation of fold change and dispersion for RNA-seq data with DESeq2. Genome Biol 15, 550 (2014).

69. Yu, G., Wang, L.-G., Han, Y. & He, Q.-Y. clusterProfiler: an R package for comparing biological themes among gene clusters. OMICS 16, 284–287 (2012).

